# Mating-compatibility genes employed as diagnostic markers to identify novel incursions of the myrtle rust pathogen *Austropuccinia psidii*

**DOI:** 10.1101/2024.02.19.580897

**Authors:** Jinghang Feng, Austin Bird, Zhenyan Luo, Rita Tam, Luc Shepherd, Lydia Murphy, Lavi Singh, Abigail Graetz, Mareike Moeller, Lilian Amorim, Nelson Sidnei Massola, M. Asaduzzaman Prodhan, Louise Shuey, Douglas Beattie, Alejandro Trujillo Gonzalez, Peri A. Tobias, Amanda Padovan, Rohan Kimber, Alistair McTaggart, Monica Kehoe, Benjamin Schwessinger, Thaís R. Boufleur

**Affiliations:** Research School of Biology, The Australian National University, Canberra, ACT 2601, Australia; DPIRD Diagnostics and Laboratory Services, Department of Primary Industries and Regional Development; 3 Baron-Hay Court, South Perth, WA 6151, Australia; Department of Plant Pathology and Nematology, Luiz de Queiroz College of Agriculture, University of São Paulo, Piracicaba, SP 13418-900, Brazil; Queensland Department of Agriculture and Fisheries, Ecosciences Precinct, 41 Boggo Rd, Dutton Park QLD 4102, Australia; Department of Health, Australian Government; Centre for Conservation Ecology and Genomics, Institute for Applied Ecology, University of Canberra, Bruce, Australian Capital Territory, Australia; School of Life and Environmental Sciences, Faculty of Science, The University of Sydney, Camperdown, New South Wales 2006, Australia; Crop Sciences, South Australian Research and Development Institute (SARDI), Waite Research Precinct, Urrbrae, SA 5064, Australia; Centre for Horticultural Science, Queensland Alliance for Agriculture and Food Innovation, The University of Queensland, Ecosciences Precinct, Dutton Park, Queensland, Australia

**Keywords:** Myrtaceae, Diagnostics, Oxford Nanopore Technologies, mating-type, Homeodomain genes

## Abstract

*Austropuccinia psidii* is the causal agent of myrtle rust in over 480 species within the family Myrtaceae. Lineages of *A. psidii* are structured by host in its native range, and some have success on new-encounter hosts. For example, the pandemic biotype has spread beyond South America, and proliferation of other lineages is an additional risk to biodiversity and industries. Efforts to manage *A. psidii* incursions, including lineage differentiation, relies on variable microsatellite markers. Testing these markers is time-consuming and complex, particularly on a large scale. We designed a novel diagnostic approach targeting the fungal mating-type *HD* (homeodomain) transcription factor locus to address these limitations. The *HD* locus (*bW1/2-*HD1 and *bE1/2-HD2)* is highly polymorphic, facilitating clear biological predictions about its inheritance from founding populations. To be considered the same lineage, all four *HD* alleles must be identical. Our lineage diagnostics relies on PCR amplification of the *HD* locus in different genotypes of *A. psidii* followed by amplicon sequencing using Oxford Nanopore Technologies (ONT) and comparative analysis. The lineage-specific assay was validated on four isolates with existing genomes, uncharacterized isolates, and directly from infected leaf material. We reconstructed *HD* alleles from amplicons and confirmed their sequence identity relative to their reference. Genealogies using *HD* alleles confirmed the variations at the *HD* loci among lineages/isolates. Our study establishes a robust diagnostic tool, for differentiating known lineages of *A. psidii* based biological predictions. This tool holds promise for detecting new pathogen incursions and can be refined for broader applications, including air-sample detection and mixed-isolate infections.

## Background

*Austropuccinia psidii*, the causal agent of myrtle rust in over 480 host species within the Myrtaceae family (Soewarto et al. 2019; Carnegie and Giblin 2021), is among the world’s top ten priority fungal species for biosecurity (Hyde et al. 2018). Its high virulence and rapid adaptability to new environments is a threat to biodiversity and industries (Chock 2020), especially in regions like Australia and New Zealand, where species of the Myrtaceae family prevail (Hyde et al. 2018). In eucalypts, for example, losses in volume due to rust severity can vary from 23% to 35% (Santos et al. 2020).

Initially described in Brazil (Winter 1884), *A. psidii* remained limited to the Americas for many decades before spreading to all continents, except Europe and Antarctica (Simpson et al. 2006). In Australia, where the pathogen was first detected in 2010 (Carnegie et al. 2010), only the pandemic biotype (group of organisms with identical genetic constitution) has been reported, and isolates belonging to exotic lineages of *A. psidii* are considered a threat to Australian natural environments and commercial native forests (DAFF 2023; Makinson et al. 2020). Disease symptoms and spore morphology are highly similar across isolates of *A. psidii* belonging to different lineages, even in cases with strong host associations (Morales et al. 2023; Boufleur et al. 2023; Ferrarezi et al. 2022). The disease is predominantly caused by the clonal stage of the pathogen and started by urediniospores. It is characterized by the initial appearance of small chlorotic spots that develop into bright orange pustules that sporulate to generate re-invective urediniospores under most conditions (Glen et al., 2007).

Early detection and diagnosis are crucial for tracking, and potentially limiting rust fungi incursions (Hussain et al. 2020). Microsatellite markers have been used to differentiate lineages of *A. psidii* (Stewart et al. 2017; Graça et al. 2013), but their application can be time-consuming and complex, particularly on a large scale. The current internationally approved assay to diagnose *A. psidii* is a species-specific qPCR (Quantitative Polymerase Chain Reaction) (IPPC 2018; Baskarathevan et al. 2016), however, the choice of gene lacks the variability needed to differentiate among pathogen lineages (Beenken 2017; Boufleur et al. 2023; Bini 2016). Therefore, there is a need to identify novel target regions that precisely diagnose different lineages of *A. psidii* for faster and precise action in a biosecurity response.

Mating in fungi is controlled through genes expressed at mating-type (*MAT*) loci (Wilson et al. 2015). In rust fungi, these are two unlinked loci, one contains pheromone precursors and receptors (*P/R*) and the other contains homeodomain (*HD*) transcription factors that are closely linked via a short DNA sequence. The *HD* locus encodes *bW-HD1* and *bE-HD2* genes, which are highly multiallelic in rust fungi (Luo et al. 2024) and many Basidiomycota (Coelho et al. 2017). These transcription factors form heterodimeric complexes between alleles and regulate cellular development during mating and the fungal life cycle (Coelho et al. 2017; Cuomo et al. 2017; Holden et al. 2023; Wilson et al. 2015). The analysis of *A. psidii* genomes confirmed physically unlinked, heterozygous *P/R* and *HD* loci, supporting that mate compatibility in this pathogen is governed by two multiallelic *HD* genes (*bW-HD1* and *bE-HD2*) and a biallelic *P/R* gene (Ferrarezi et al. 2022).

The aim of this study was to develop a highly sensitive assay for the detection and identification of *A. psidii* lineages distinct from the pandemic biotype. This assay can be used for monitoring existing incursions/outbreaks, and to help prevent and limit further incursions of exotic lineages. Here we introduce novel primers designed to target the mating-type *HD* locus of *A. psidii*. These primers are designed to be used in combination with long-read sequencing such as those facilitated by Oxford Nanopore Technologies (ONT).□□□

## Material and Methods

### HD locus identification and primer design

The *HD* locus of three *A. psidii* lineages was identified on complete dikaryotic genome assemblies, including Brazilian isolates belonging to two different lineages MF-1 (from *Eucalyptus grandis*) (PRJNA215767, GCA_000469055.2) and LFNJM1 (unpublished data) from *Syzygium jambos* (Boufleur et al. 2023), and APG1 from *Psidium guajava* 2/28/2024 8:02:00 AM (unpublished data), along with the Au3 isolate that belongs to the pandemic biotype (Au3_v2) (PRJNA810573, GCA_023105745.1, GCA_023105775.1) (Edwards et al. 2022), and the South African isolate Apsidii_AM, that belongs to the South African Biotype (PRJNA480390, GCA_003724095.1) (McTaggart et al. 2018). The *HD* loci containing regions were identified with BLASTx (v.2.15.0) in combination with annotated *bW-HD1* and *bE-HD2 A. psidii* genes, as described by Ferrarezi et al., (2022). As expected for the *HD* locus in dikaryotic genome assemblies, two alleles of each *bW-HD1* and *bE-HD2* gene were retrieved. The alleles of each gene (*bW-HD1* and *bE-HD2*) were aligned separately with MACSE (v.2.07) (Ranwez et al. 2018) and two Bayesian inference genealogy trees were generated with Mr. Bayes v.3.2.6 (Huelsenbeck and Ronquist 2001).

Primers were designed manually based on a multiple sequence alignment of contigs containing the *HD* alleles of the pandemic Au3_v2 isolate, the South African isolate Apsidii_AM, and the Brazilian isolate MF-1. Two pairs of degenerate primers were designed to amplify *bW-HD1* and *bE-HD2* individually, and the combination of the most forward and the most reverse primers was used to amplify the full-length *HD* locus (Table 1). Regions that were fully or nearly fully conserved having a maximum of two single nucleotide differences between all *HD* loci alleles were selected to allow amplification of all alleles at the same time. The expected amplicon size was ∼1600 bp and ∼1400 bp for each *bW-HD* and *bE-HD,* and ∼3000 bp for the complete *HD* locus.

**Table 1.**
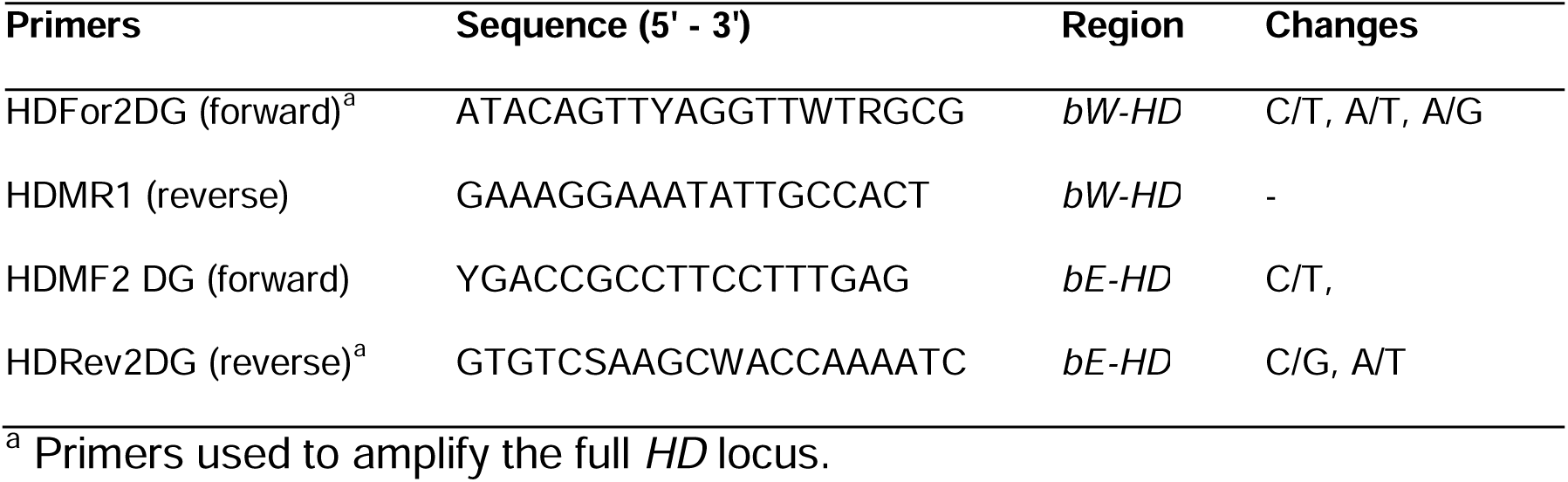
List of primers used in the present study.

### DNA extraction and HD locus amplification

The designed primers were tested with diverse samples, including four positive controls of *A. psidii*, ten field samples and three non-target rust species (Table 2). Genomic DNA (gDNA) was extracted directly from *A. psidii* urediniospores, from infected leaf material or from urediniospores of non-target rust species (Table 2) with the DNeasy Plant mini kit (Qiagen) according to the manufacturer’s instructions. The integrity and quality of the DNA was measured with a Nanodrop spectrophotometer (Thermo Fisher Scientific) and checked by agarose gel (0.8%) electrophoresis stained with SYBR Safe (Thermo Fisher Scientific). The DNA concentration was determinate using the Qubit 4 (Thermo Fisher Scientific), and all samples were adjusted to 25 ng/µL for further studies.

**Table 2.**
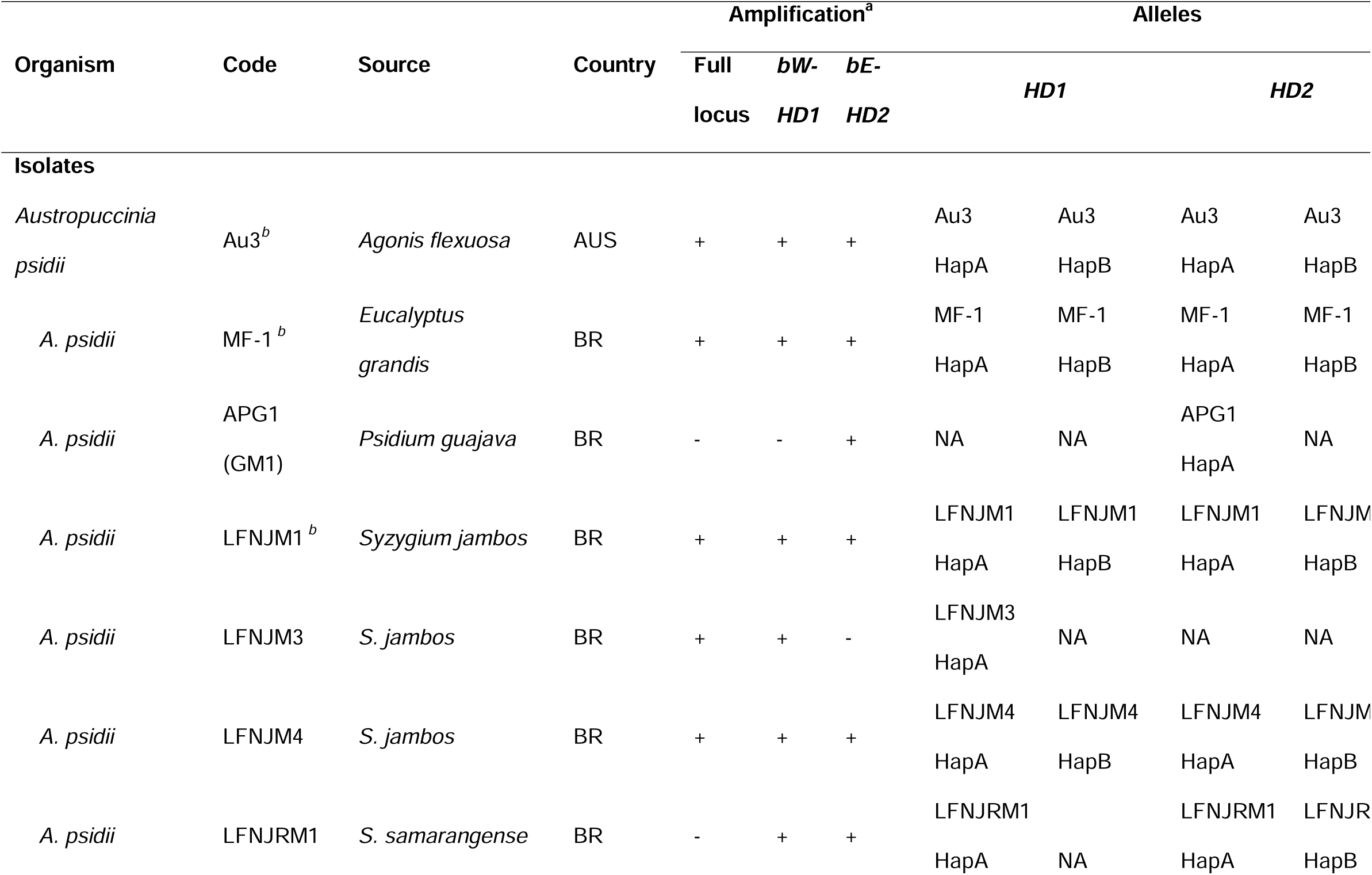
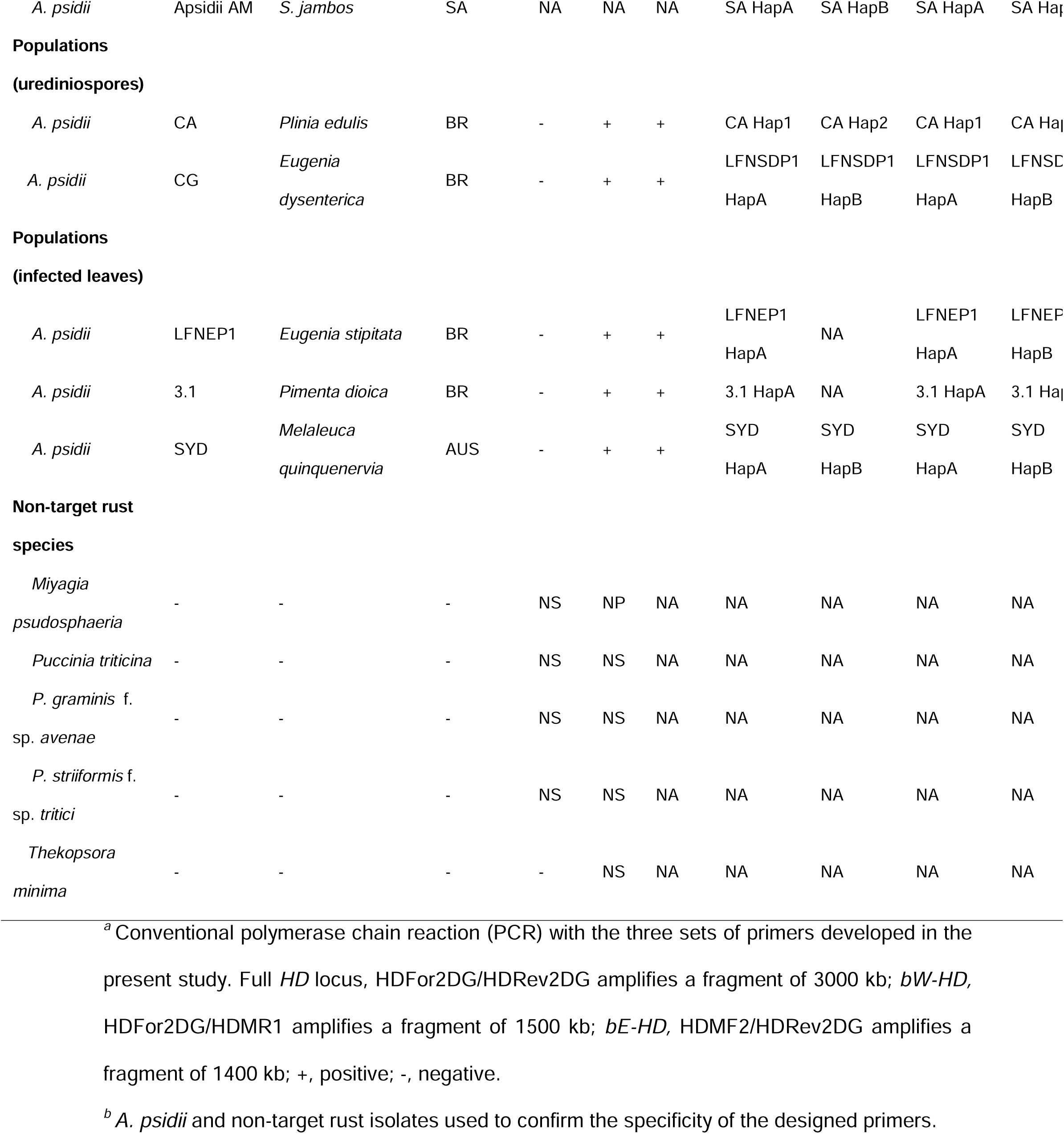
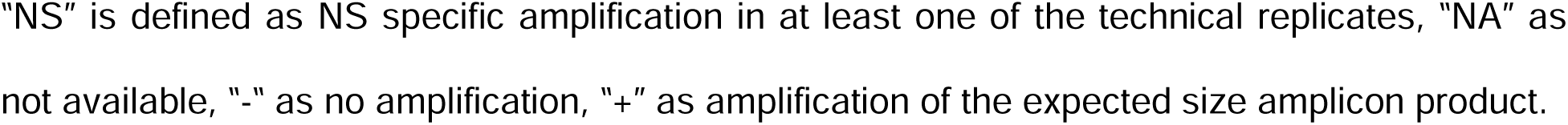
Description of isolates and populations and of *A. psidii* and non-target rusts used in this study.

In the first round of PCR tests, the aim was to evaluate the amplification of (non-) target sequences by the designed primers followed by sequence analysis using *A. psidii* and three non-target rust isolates (Table 2, Figure 1A-B). PCR was performed on a Mastecycler nexus X2 thermal cycler (Eppendorf). The reaction mixture, with a final volume of 25 µL, included 5 µL of 5X reaction buffer, 0.5 µL of dNTPs [10 mM], 1.25 µL of each primer [10 µM], 0.25 µL of Q5 High-Fidelity DNA Polymerase (NEB), 14.75 uL of Nuclease Free Water (NFW) and 2 µL (up to 50 ng) of template DNA. The PCR amplification had an initial denaturation step at 98 °C for 30 s, followed by 35 cycles of denaturation at 98 °C for 30 s, annealing at 58 °C for 30 s and extension at 72 °C for 30 s, with a final extension step at a 72 °C for 2 min. Specificity tests were run in duplicate, and PCR products were visualised on 2% agarose gel stained with SYBR safe (Thermo Fisher Scientific).

**Figure 1:**
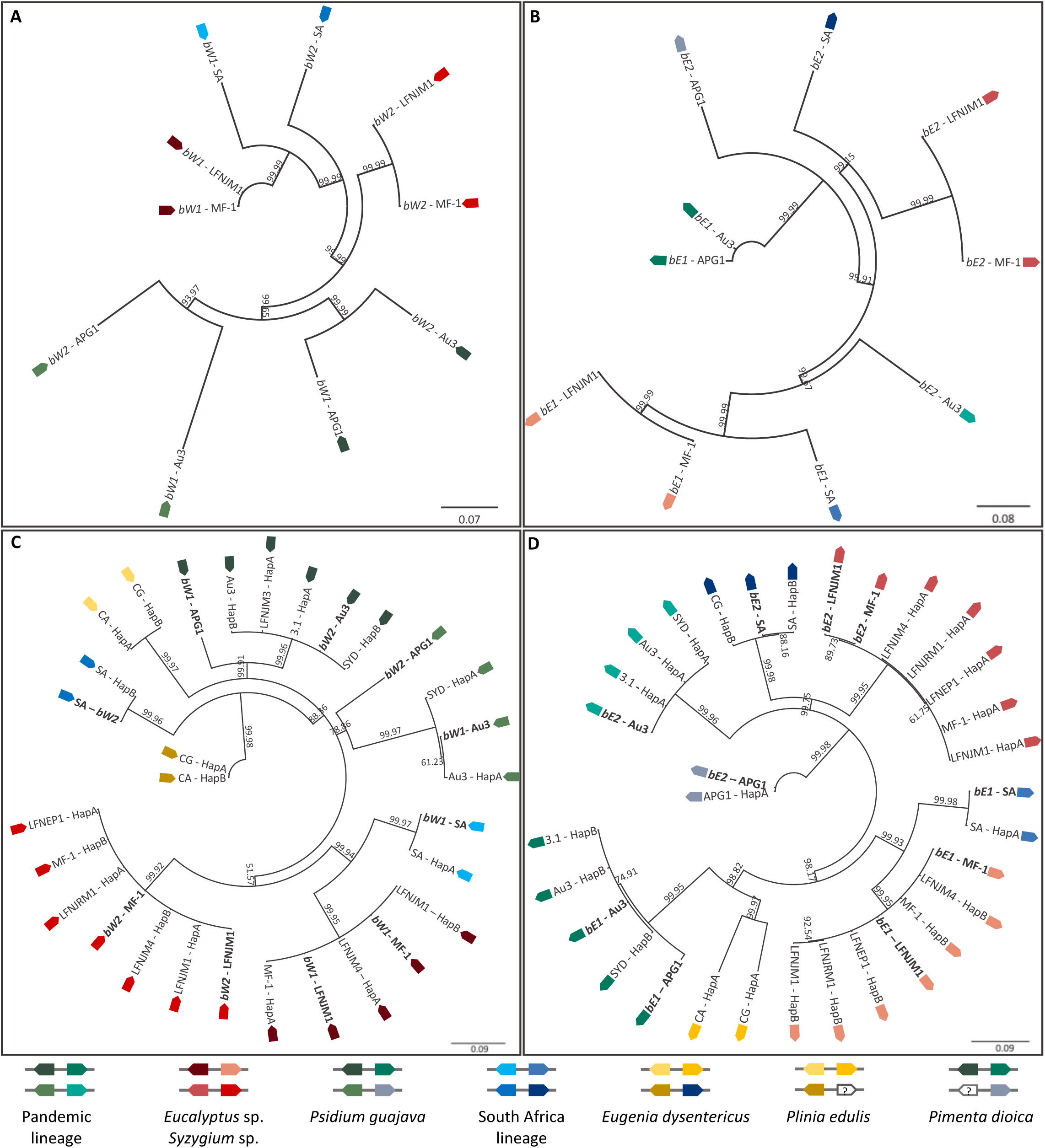
Bayesian inference genealogical trees reconstructed from *HD* (homeodomain) genes of *Austropuccinia psidii*. Genealogical trees reconstructed from the alignment of two *bW-HD1* **(A, C)** or two *bE-HD2* **(B, D)** alleles per dikaryotic genome assembly of reference isolates **(A-B),** and with consensus sequences for each allele *de novo* reconstructed from ONT long-read amplicons of *bW-HD1* and *bE-HD2* (C-D). Lineage identification is indicated by the combination of different colors in the bottom. Reference sequences are indicated in bold (C-D). Bayesian posterior probabilities are shown on the nodes.

In the second round of PCR tests, the performance of the primers was assessed against single-spore isolates and field samples (urediniospores and infected leaves) collected in Brazil and Australia (Table 2). Identical reaction conditions, as described above, were used to amplify the *bW-HD* and *bE-HD* loci individually or the full-length *HD locus* (Table 2).

### ONT sequencing

For Oxford Nanopore sequencing, libraries were generated following manufacturer instructions for V14 Ligation Sequencing of amplicons (Native Barcoding Kit V14 96 - SQK-NBD114.96) with modifications as follows. An initial bead clean was performed using 1.2 x of 2 % Sera-Mag beads to purify the PCR product and 200 fmol of clean DNA carried through to End-prep reaction. The End-prep reaction was incubated at 20 °C and then 65 °C for 15 min to maximize yield. One µL of the end-prepped DNA was amplicons were barcoded with 1 µL of Nanopore Native Barcode using 5 µL Blunt/TA ligase master mix (NEB) in total reaction volume of 10 µL for 20 min at 20 °C. The reaction was stopped by adding 1 µL of EDTA to each ligation reaction. The individually barcoded PCR amplicons were pooled, bead cleaned with 0.6 x volume 2% Sera-Mag beads and two washes of 70% ethanol. The pool of barcoded PCR amplicons was eluted in 21 µL of nuclease-free water. Library preparation was completed according to the manufacturers protocol. Twenty fmol of the barcoded library was loaded on a MinION R10.4.1 Flowcell (FLO-MIN114) and sequencing was ran using a MinION. Basecalling was performed with Guppy v. 6.4.2 Super High Accuracy mode. All long-read amplicon datasets were deposited to Zenodo (https://doi.org/10.5281/zenodo.10656657).

### De novo reconstruction of HD loci and genealogies

A two-step filtering process was implemented on the base-called sequences. In the initial step, sequencing reads were filtered based on their Phred quality scores, with reads having a mean quality score below 15 being removed. This ensured that the remaining reads had an average per base accuracy of ≥ 97%. The second filtering step involved selecting sequencing reads with lengths around the expected amplicon length for each *HD* gene region: 1500-1800 bp for *bW-HD1* and 1300-1500 bp for *bE-HD2*. The VSEARCH clustering algorithm (Rognes et al. 2016) was applied for quality control of sequencing reads of each sample, using global similarity during clustering. It is important to note that the VSEARCH clustering algorithm could not recognize sequences of the same gene in the opposite direction. Hence, we obtained two clusters for each *bW-HD1* and *bE-HD2* alleles in the sample along with two consensus sequences - one forward and the other reverse. The analysis code is available on Github (https://github.com/TheRainInSpain/Lineage-Specific-Marker.git). And the dataset is available on (https://doi.org/10.5281/zenodo.10656657). Geneious software (v.2023.2.1) was used to visualize the forward read consensus sequences.

The reconstructed amplicon and reference sequences were aligned (Table 2) with MUSCLE (v.5.1) (Edgar 2004), and simple genealogical trees were reconstructed using the Geneious tree builder (v.2023.2.1).

## Results

### Specificity of the diagnostic assay achieved through HD loci amplification and ONT sequencing

During the initial phase of primer testing, *HD* PCRs were performed on positive controls of *A. psidii* (Au3, MF-1, APG1 and LFNJM1) and negative controls with non-target rust species (*Miyagia psudosphaeria, Puccinia striiformis* f. sp. *tritici, P. graminis*, *P. triticina, Thekopsora minima*). In addition, we focused on the individual PCR amplicons (*bW-HD* and *bE-HD*) because the amplification of the full-length locus was not robust enough across samples and technical replicates. The sizes of the amplicons were as expected in the positive control samples, being of ∼1600 bp for *bW-HD*, ∼1400 bp for *bE-HD*, and ∼3000 kb for the full HD locus (Table 2). In addition, we observed bands of variable sizes in some of our technical repeats of non-target species. All samples were sequenced with our ONT amplicon sequencing workflow because our assay does not rely exclusively on the PCR amplification product but requires that the amplified sequences match the *A. psidii HD* sequences in a genealogical framework.

None of the non-target species amplicon sequences matched the full-length *A. psidii HD* sequences. For each *A. psidii* isolate, four *HD* alleles (2x *bW-HD1* and 2x *bE-HD2*) were reconstructed based on ONT sequencing results, as expected for dikaryotic organisms. Isolates were considered identical if their four *HD* alleles were identical or carried minor non-functional variation (for example synonymous variation, variation outside the variable domain or within introns). The applicability of the lineage diagnostic test was confirmed by comparing the *de novo* reconstructed consensus sequences derived from ONT amplicon sequencing with the *HD* amplicon sequences derived from the reference genomes (APG1, MF-1, LFNJM1, Au3 and Apsidii_AM). All *de novo* reconstructed ONT amplicon sequences clearly grouped with their respective *HD* alleles obtained from reference genomes (Table 2, Figure 1A-B). Moreover, the reference trees for *bW-HD1* and *bE-HD2* revealed that the Brazilian isolates and the South African isolate carry clearly distinct alleles when compared against the pandemic isolate (Figure 1A-B), corroborating variations previously described using microsatellite markers (Stewart et al. 2017; Roux et al. 2016; Graça et al. 2013).

### Primers targeting the *HD* region distinguished different lineages of *A. psidii*

The designed primers successfully amplified individual *HD* loci of DNA extracted from different sources of field samples, including urediniospores and infected leaf material. As observed previously, the full *HD* locus amplification was not possible for the LFNJRM1 isolate and the field samples (Table 2). The reconstructed amplicon and reference sequences were aligned and revealed a pairwise identity of 67.9% to 100% for *bW-HD1* and 76.9% and to 100% for *bE-HD2* alleles across the analyzed sample isolates. Our results revealed that an isolate collected from field samples in Australia in 2022 (SYD) belong to the same lineage as the pandemic lineage because all four *HD* alleles clearly grouped with those derived from the pandemic reference isolate, while isolates collected from field samples from Brazil, belong to different lineages having at least two different *HD* alleles (Figure 1C-D).

## Discussion

Traditionally, *A. psidii* lineages have been identified using microsatellite markers (Stewart et al. 2017; Graça et al. 2013; Roux et al. 2016; Sandhu et al. 2016). In this study, a diagnostic assay targeting the *HD* locus of *A. psidii*, coupled with ONT sequencing was developed. This method successfully amplified diverse copies of the *HD* locus from *A. psidii* isolates collected in Australia and Brazil and is predicted to work for the South African isolate.

The genealogical tree of *de novo* reconstructed amplicons confirmed the biological expectation of lineage specific variation among isolates originating from single spores, field samples from different hosts, and distinct geographic locations. The isolate collected from field samples in Australia had *HD* alleles that matched the pandemic lineage, whereas in Brazil, the center of origin of the disease, there was a strong association of *HD* allele status with their original host species, as previously observed (Morales et al. 2023; Graça et al. 2013; Stewart et al. 2017). Overall, the high allelic diversity at the *HD* locus in our samples is in line with the previously identified diversity observed at this locus in rust fungi including *A. psidii* and *Puccinia* spp. (Ferrarezi et al. 2022; Holden et al. 2023; Luo et al., 2024).

Individual *HD* genes were amplified in field samples for which the entire *HD* locus could not be amplified by PCR. In the future, our primers and/or amplification conditions can be further improved to amplify the entire locus of natural *A. psidii* populations within its center of diversity, as it is easier in a Biosecurity response to amplify one long region than two. In addition, we observed off target PCR amplification in some of our technical replicates of non-target species, yet none of these sequences match the *A. psidii HD* sequence. Improved primers or nested PCRs could improve PCR specificity increasing sensitivity and throughput when applied under routine testing conditions.

To date, the non-pandemic isolates shared at most one out of four *HD* alleles with the pandemic isolate (Au_3). This finding shows non-pandemic isolates can be differentiated from the pandemic lineage based on their *HD* alleles. This protocol serves as a valuable tool in detecting new incursions of the pathogen in regions where a single lineage is present as for example in Australia where only a single incursion of the pandemic biotype has been reported to date. Further, comparing *HD* allele identity with known reference sequences could link novel incursions with related populations in source regions. This could help in identifying risks in import pathways of this exotic pathogen and improve risk mitigation strategies. In addition, the assay could potentially detect recombination between populations if purified single pustule isolates were analyzed. In the future, we anticipate that the diagnostic test can be refined to detect urediniospores of *A. psidii* from complex samples derived from air-sampling or mixed infections to enable structured targeted surveillance of this pathogen on the ground.

## Acknowledgments

The authors thank Danièle Giblot-Ducray and Kelly Hill for the critical feedback and support.

